## Supplemental Table S1 for "The Kinetochore Protein KNL-1 Regulates the Actin Cytoskeleton to Control Dendrite Branching"

**Table S1: *C. elegans* strain list**

| **REAGENT OR RESOURCE** | **SOURCE** | **IDENTIFIER** |
| --- | --- | --- |
| **Experimental Model: *C. elegans* Strains** |  |  |
| *C. elegans N2 Bristol* | Caenorhabditis Genetics Center | N2 |
| *dhaSi76 [oxTi177; pDC731; Pdes-2::mScarlet-I::PH::unc-54::3'UTR; cb-unc-119(+)]IV* | This study | DKC410 |
| *dhaSi145 [oxTi185; pDC793; Pdes‑2::TIR1::F2A::mTagBFP::AID::NLS::tbb-2::3'UTR; cb-unc-119(+)]I; dhaSi89 [pDC747; Punc-86::TIR-1::unc-54::3'UTR; cb-unc-119(+)]II; dhaSi76 [oxTi177; pDC731; Pdes-2::mScarlet-I::PH::unc-54::3'UTR; cb-unc-119(+)]IV* | This study | DKC815 |
| *dhaEx1[Pdes-2::knl-1::GFP::unc-54::3'UTR]; dhaSi76[oxTi177; pDC731; Pdes-2::mScarlet-I::PH::unc-54::3'UTR; cb-unc-119(+)]IV* | This study | DKC822 |
| *dhaEx3[Pdes-2::GFP::unc-54::3'UTR]; dhaSi76[oxTi177; pDC731; Pdes-2::mScarlet-I::PH::unc-54::3'UTR; cb-unc-119(+)]IV* | This study | DKC824 |
| *dhaSi145 [oxTi185; pDC793; Pdes‑2::TIR1::F2A::mTagBFP::AID::NLS::tbb-2::3'UTR; cb-unc-119(+)]I; dhaSi89 [pDC747; Punc-86::TIR-1::unc-54::3'UTR; cb-unc-119(+)]II; knl-1 (dha122 [AID:: mTagBFP::knl-1])III; dhaSi76 [oxTi177; pDC731; Pdes-2::mScarlet-I::PH::unc-54::3'UTR; cb-unc-119(+)]IV* | This study | DKC849 |
| *dhaSi145 [oxTi185; pDC793; Pdes‑2::TIR1::F2A::mTagBFP::AID::NLS::tbb-2::3'UTR; cb-unc-119(+)]I; dhaSi89 [pDC747; Punc-86::TIR-1::unc-54::3'UTR; cb-unc-119(+)]II; ndc-80 (dha148 [ndc-80::AID::dGFP::3XFLAG]); dhaSi76 [oxTi177; pDC731; Pdes-2::mScarlet-I::PH::unc-54::3'UTR; cb-unc-119(+)]IV* | This study | DKC860 |
| *dhaSi207 [pDC1020; Punc-86::splitGFP1-10::tbb-2::3'UTR; cb‑unc‑119(+)]II; knl-1(dha130 [HA::7XGFP11::knl-1])III; dhaSi76 [oxTi177; pDC731; Pdes-2::mScarlet-I-PH::unc-54::3'UTR; cb‑unc‑119(+)]IV* | This study | DKC870 |
| *dhaSi207 [pDC1020; Punc-86::splitGFP1-10::tbb-2::3'UTR; cb-unc-119(+)]II; knl-1(dha130 [HA::7XGFP11::knl-1])III; dhaSi249 [oxTi365; pDC732; Punc-86::mScarlet-I::PH::unc-54::3'UTR]V* | This study | DKC872 |
| *dhaSi145 [oxTi185; pDC793; Pdes‑2::TIR1::F2A::mTagBFP::AID::NLS::tbb-2::3'UTR; cb-unc-119(+)]I; dhaSi89 [pDC747; Punc-86::TIR-1::unc-54::3'UTR; cb-unc-119(+)]II; dhaSi76 [oxTi177; pDC731; Pdes-2::mScarlet-I::PH::unc-54::3'UTR; cb-unc-119(+)]IV; knl-3(du2 [AID::knl-3])V* | This study | DKC911 |
| *dhaSi145 [oxTi185; pDC793; Pdes-2::TIR1::F2A::mTagBFP::AID::NLS::tbb-2::3’UTR; cb-unc-119(+)]I; dhaSi89 [pDC747; Punc-86::TIR-1::unc-54::3’UTR; cb-unc-119(+)]II; knl-1(dha122 [AID::mTagBFP::knl-1])III; dhaSi76 [oxTi177; pDC731; Pdes-2::mScarlet-I::PH::unc-54::3'UTR; cb-unc-119(+)]IV; dhaSi241 [oxTi365; pDC978; Pdes-2::GFP::tba-1::unc-54::3'UTR; cb-unc-119(+)]V* | This study | DKC916 |
| *dhaSi145 [oxTi185; pDC793; Pdes-2::TIR1::F2A::mTagBFP::AID::NLS::tbb-2::3’UTR; cb-unc-119(+)]I; dhaSi89 [pDC747; Punc-86::TIR-1::unc-54::3’UTR; cb-unc-119(+)]II; dhaSi76 [oxTi177; pDC731; Pdes-2::mScarlet-I::PH::unc-54::3'UTR; cb-unc-119(+)]IV; dhaSi241[oxTi365; pDC978; Pdes-2::GFP::tba-1::unc-54::3'UTR; cb-unc-119(+)]V* | This study | DKC917 |
| *dma-1(dha155; DEL)I* | This study | DKC919 |
| *dhaSi145 [oxTi185; pDC793; Pdes‑2::TIR1::F2A::mTagBFP::AID::NLS::tbb-2::3’UTR; cb-unc-119(+)]I; dhaSi89 [pDC747; Punc-86::TIR-1::unc-54::3’UTR; cb-unc-119(+)]II; hrtSi17 [Pdes-2::mKate2::rab-3 LG]IV* | This study | DKC1001 |
| *dhaSi145 [oxTi185; pDC793; Pdes‑2::TIR1::F2A::mTagBFP::AID::NLS::tbb-2::3’UTR; cb-unc-119(+)]I; dhaSi89 [pDC747; Punc-86::TIR-1::unc-54::3’UTR; cb-unc-119(+)]II; knl-1(dha122 [AID::mTagBFP::knl-1])III; hrtSi17 [Pdes-2::mKate2::rab-3 LG]IV* | This study | DKC1002 |
| *dhaSi145 [oxTi185; pDC793; Pdes-2::TIR1::F2A::mTagBFP::AID::NLS::tbb-2::3’UTR; cb-unc-119(+)]I; dhaSi89 [pDC747; Punc-86::TIR-1::unc-54::3'UTR; cb-unc-119(+)]II; dhaSi243 [pDC929; oxTi177; Pdes-2::Lifeact::mKate2::tbb-2::3’UTR; cb-unc-119(+)]IV* | This study | DKC1035 |
| *dhaSi145 [oxTi185; pDC793; Pdes-2::TIR1::F2A::mTagBFP::AID::NLS::tbb-2::3’UTR; cb-unc-119(+)]I; dhaSi89 [pDC747; Punc-86::TIR-1::unc-54::3’UTR; cb-unc-119(+)]II; knl-1(dha122 [AID::mTagBFP::knl-1])III; dhaSi243 [pDC929; oxTi177; Pdes-2::Lifeact::mKate2::tbb-2::3’UTR; cb-unc-119(+)]IV* | This study | DKC1057 |
| *dhaSi145 [oxTi185; pDC793; Pdes‑2::TIR1::F2A::mTagBFP::AID::NLS::tbb-2::3’UTR; cb-unc-119(+)]I; dhaSi89 [pDC747; Punc-86::TIR-1::unc-54::3'UTR; cb-unc-119(+)]II; knl-1(dha122 [AID::mTagBFP::knl-1])III; dhaSi76 [oxTi177; pDC731; Pdes-2::mScarlet-I::PH::unc-54::3'UTR; cb-unc-119(+)]IV; dhaSi269 [oxTi365; pDC1106; Pser‑2prom3::ebp-2::GFP::tbb-2::3'UTR; cb-unc-119(+)]V* | This study | DKC1072 |
| *dhaSi145 [oxTi185; pDC793; Pdes‑2::TIR1::F2A::mTagBFP::AID::NLS::tbb-2::3’UTR; cb-unc-119(+)]I; dhaSi89 [pDC747; Punc-86::TIR-1::unc-54::3’UTR; cb-unc-119(+)]II; dhaSi76 [oxTi177; pDC731; Pdes-2::mScarlet-I::unc-54::3'UTR; cb-unc-119(+)]IV; dhaSi269 [oxTi365; pDC1106; Pser-2prom3::ebp-2::GFP::tbb-2::3'UTR; cb-unc119(+)]V* | This study | DKC1073 |
| *dhaSi145 [oxTi185; pDC793; Pdes-2::TIR1::F2A::mTagBFP::AID::NLS::tbb-2::3’UTR; cb-unc-119(+)]I; dhaSi89 [pDC747; Punc-86::TIR-1::unc-54::3’UTR; cb-unc-119(+)]II; knl-1(dha122 [AID::mTagBFP::knl-1])III; dhaSi272 [oxTi365; pDC1122; Pser2-prom3::dma-1::mKate2::unc-54::3'UTR;cb-unc-119(+)]V* | This study | DKC1075 |
| *dhaSi145 [oxTi185; pDC793; Pdes-2::TIR1::F2A::mTagBFP::AID::NLS::tbb-2::3’UTR; cb-unc-119(+)]I; dhaSi89 [pDC747; Punc-86::TIR-1::unc-54::3’UTR; cb-unc-119(+)]II; dhaSi272 [oxTi365; pDC1122; Pser-2prom3::dma-1::mKate2::unc-54::3'UTR; cb-unc-119(+)]V* | This study | DKC1076 |
| *dhaSi145 [oxTi185; pDC793; Pdes‑2::TIR1::F2A::mTagBFP::AID::NLS::tbb-2::3’UTR; cb-unc-119(+)]I; tba-1(dha149[HA::GFP11::tba-1])I; dhaSi89 [pDC747; Punc-86::TIR-1::unc-54::3’UTR; cb-unc-119(+)]II; dhaSi76[oxTi177; pDC731; Pdes-2::mScarlet-I::PH::unc-54::3'UTR; cb-unc-119(+)]IV; dhaSi207 [oxTi365; pDC1020; Punc-86::splitGFP1-10::tbb-2::3'UTR; cb-unc-119(+)]V* | This study | DKC1086 |
| *dhaSi145 [oxTi185; pDC793; Pdes‑2::TIR1::F2A::mTagBFP::AID::NLS::tbb-2::3'UTR; cb-unc-119(+)]I; dhaSi89 [pDC747; Punc-86::TIR-1::unc-54::3'UTR; cb-unc-119(+)]II; dhaSi243 [oxTi177; pDC929; Pdes‑2::Lifeact::mKate2::tbb-2::3’UTR; cb-unc-119(+)]IV; dhaSi45 [oxTi365; pDC675; Pdes-2::mNeonGreen::PH::unc-54::3'UTR; cb-unc-119(+)]V* | This study | DKC1087 |
| *dhaSi145 [oxTi185; pDC793; Pdes‑2::TIR1::F2A::mTagBFP::AID::NLS::tbb-2::3'UTR; cb-unc-119(+)]I; dhaSi89 [pDC747; Punc-86::TIR-1::unc-54::3'UTR; cb-unc-119(+)]II; knl-1(dha122[AID::mTagBFP::knl-1])III; dhaSi45 [oxTi365; pDC675; Pdes-2::mNeonGreen::PH::unc-54::3'UTR; cb-unc-119(+)]V* | This study | DKC1089 |
| *dhaSi145 [oxTi185; pDC793; Pdes-2::TIR1::F2A::mTagBFP::AID::NLS::tbb-2::3’UTR; cb-unc-119(+)]I; tba-1(dha149[HA::GFP11::tba-1])I; dhaSi89 [pDC747; Punc-86::TIR-1::unc-54::3'UTR; cb-unc-119(+)]II; knl-1(dha122 [AID::mTagBFP::knl-1])III; dhaSi76[oxTi177; pDC731; Pdes-2::mScarlet-I::PH::unc-54::3'UTR; cb-unc-119(+)]IV; dhaSi207 [oxTi365; pDC1020; Punc-86::splitGFP1-10::tbb-2::3'UTR; cb-unc‑119(+)]V* | This study | DKC1090 |
| *dhaSi145 [oxTi185; pDC793; Pdes‑2::TIR1::F2A::mTagBFP::AID::NLS--tbb-23'UTR; cb-unc-119(+)]I; dhaSi89 [pDC747; Punc-86::TIR-1::unc-54::3’UTR; cb-unc-119(+)]II; knl-1(dha122 [AID::mTagBFP::knl-1])III; dhaSi243 [pDC929; oxTi177; Pdes‑2::Lifeact::mKate2::tbb-2::3’UTR; cb-unc-119(+)]IV; dhaSi45 [oxTi365; pDC675; Pdes-2::mNeonGreen::PH::unc-54::3'UTR; cb-unc-119(+)]V* | This study | DKC1113 |
| *dhaSi145 [oxTi185; pDC793; Pdes-2::TIR1::F2A::mTagBFP::AID::NLS::tbb-2::3’UTR; cb-unc-119(+)]I; dhaSi89 [pDC747; Punc-86::TIR-1::unc-54::3'UTR; cb-unc-119(+)]II; knl-1(dha122 [AID::mTagBFP::knl-1])III; dhaSi237[oxTi365; pDC1061;Pser-2prom3::mScarlet‑I::LGG1::unc-54::3'UTR; cb-unc-119(+)]V* | This study | DKC1126 |
| *dhaSi145 [oxTi185; pDC793; Pdes-2::TIR1::F2A::mTagBFP::AID::NLS::tbb-2::3’UTR; cb-unc-119(+)]I; dhaSi89 [pDC747; Punc-86::TIR-1::unc-54::3'UTR; cb-unc-119(+)]II; dhaSi237 [oxTi365; pDC1061;Pser-2prom3::mScarlet‑I::LGG1::unc-54::3'UTR; cb-unc-119(+)]V* | This study | DKC1127 |
| *dma-1 (dha155 [DEL])I; dhaSi89 [pDC747; Punc-86::TIR-1::unc-54::3'UTR; cb-unc-119(+)]II; mec-4(dha181[mec-4(e1611 allele mutant)])X* | This study | DKC1128 |
| *dhaSi145 [oxTi185; pDC793; Pdes-2::TIR1::F2A::mTagBFP::AID::NLS::tbb-2::3'UTR; cb-unc-119(+)]I; dhaSi89 [pDC747; Punc-86::TIR-1::unc-54::3'UTR; cb-unc-119(+)]II; knl-1(dha122 [AID::mTagBFP::knl-1])III; dhaSi53 [oxTi177; pDC675; Pdes-2::mNeonGreen::PH::unc-54::3'UTR; cb-unc-119(+)]IV; dhaSi237 [oxTi365; pDC1061;Pser-2prom3::mScarlet‑I::LGG1::unc-54::3'UTR; cb-unc-119(+)]V* | This study | DKC1131 |
| *dhaSi145 [oxTi185; pDC793; Pdes-2::TIR1::F2A::mTagBFP::AID::NLS::tbb-2::3'UTR; cb-unc-119(+)]I; dhaSi89 [pDC747; Punc-86::TIR-1::unc-54::3'UTR; cb-unc-119(+)]II; dhaSi53 [oxTi177; pDC675; Pdes-2::mNeonGreen::PH::unc-54::3'UTR; cb-unc-119 (+)]IV; dhaSi237[oxTi365; pDC1061;Pser-2prom3::mScarlet‑I::LGG1::unc-54::3'UTR; cb-unc-119(+)]V* | This study | DKC1132 |
| *dhaSi89 [pDC747; Punc-86::TIR-1::unc-54::3’UTR; cb-unc-119(+)]II; knl-1(dha130 [HA::7XGFP11::knl-1])III; dhaSi76 [oxTi177; pDC731; Pdes-2::mScarlet-I::PH::unc-54::3'UTR; cb-unc-119(+)]IV; dhaSi207 [oxTi365; pDC1020; Punc-86::splitGFP1-10::tbb-2::3'UTR; cb-unc-119(+)]V* | This study | DKC1139 |
| *dhaSi89 [pDC747; Punc-86::TIR-1::unc-54::3’UTR; cb-unc-119(+)]II; knl-1(dha185 [HA::7XGFP11::knl-1::AID])III; dhaSi76 [oxTi177; pDC731; Pdes-2::mScarlet-I-PH::unc-54::3'UTR; cb-unc-119(+)]IV; dhaSi207 [oxTi365; pDC1020;Punc-86::splitGFP1-10::tbb-2::3'UTR; cb-unc-119(+)]V* | This study | DKC1151 |
| *dhaSi89 [pDC747; Punc-86::TIR-1::unc-54::3'UTR; cb-unc-119(+)]II; mec-4(dha181 [mec-4(e1611 allele mutant)])X* | This study | DKC1158 |
| *dhaSi277 [pDC1080; Pdes-2::MYR::KNL-1::mTagBFP::unc-54::3'UTR; cb-unc-119(+)]II; dhaSi243 [pDC929; oxTi177; Pdes‑2::Lifeact::mKate2::tbb-2::3’UTR; cb-unc-119(+)]IV; dhaSi45 [oxTi365; pDC675; Pdes-2::mNeonGreen::PH::unc-54::3'UTR; cb-unc-119(+)]V* | This study | DKC1251 |
| *dhaSi307 [pDC1079; Pdes-2::MYR::mTagBFP::unc-54::3'UTR; cb-unc-119(+)]II; dhaSi243 [pDC929; oxTi177; Pdes‑2::Lifeact::mKate2::tbb-2::3’UTR; cb-unc-119(+)]IV; dhaSi45 [oxTi365; pDC675; Pdes-2::mNeonGreen::PH::unc-54::3'UTR; cb-unc-119(+)]V* | This study | DKC1253 |
| *dhaSi311 [pDC1081; Pdes-2::MYR::KNL-1(1-505) ::mTagBFP::unc-54::3'UTR; cb-unc-119(+)]II; dhaSi243 [pDC929; oxTi177; Pdes‑2::Lifeact::mKate2::tbb-2::3’UTR; cb-unc-119(+)]IV; dhaSi45 [oxTi365; pDC675; Pdes-2::mNeonGreen::PH::unc-54::3'UTR; cb-unc-119(+)]V* | This study | DKC1299 |
| *dhaSi312 [pDC1082; Pdes-2::MYR::KNL-1(505-1010)::mTagBFP::unc-54::3'UTR; cb-unc-119(+)]II; dhaSi243 [pDC929; oxTi177; Pdes‑2::Lifeact::mKate2::tbb-2::3’UTR; cb-unc-119(+)]IV; dhaSi45 [oxTi365; pDC675; Pdes-2::mNeonGreen::PH::unc-54::3'UTR; cb-unc-119(+)]V* | This study | DKC1300 |
| *dhaSi145 [oxTi185; pDC793; Pdes‑2::TIR1::F2A::mTagBFP::AID::NLS::tbb-2::3'UTR; cb-unc-119(+)]I; dhaSi89 [pDC747; Punc-86::TIR-1::unc-54::3'UTR; cb-unc-119(+)]II; cyk-1 (dha230 [cyk-1::AID])III; dhaSi76 [oxTi177; pDC731; Pdes-2::mScarlet-I::PH::unc-54::3'UTR; cb-unc-119(+)]IV* | This study | DKC1406 |
| *dhaSi145 [oxTi185; pDC793; Pdes‑2::TIR1::F2A::mTagBFP::AID::NLS::tbb-2::3'UTR; cb-unc-119(+)]I; dhaSi89 [pDC747; Punc-86::TIR-1::unc-54::3'UTR; cb-unc-119(+)]II; knl-1 (dha122 [AID:: mTagBFP::knl-1]); cyk-1 (dha230 [cyk-1::AID])III; dhaSi76 [oxTi177; pDC731; Pdes-2::mScarlet-I::PH::unc-54::3'UTR; cb-unc-119(+)]IV* | This study | DKC1408 |
| *dhaSi338 [pDC1258; Pdes-2::MYR::KNL-1(MELA)::mTagBFP::unc::54::3'UTR; cb-unc-119(+)]II; dhaSi243 [pDC929; oxTi177; Pdes‑2::Lifeact::mKate2::tbb-2::3’UTR; cb-unc-119(+)]IV; dhaSi45 [oxTi365; pDC675; Pdes-2::mNeonGreen::PH::unc-54::3'UTR; cb-unc-119(+)]V* | This study | DKC1425 |
| *dhaSi145 [oxTi185; pDC793; Pdes‑2::TIR1::F2A::mTagBFP::AID::NLS::tbb-2::3'UTR; cb-unc-119(+)]I; dhaSi89 [pDC747; Punc-86::TIR-1::unc-54::3'UTR; cb-unc-119(+)]II; dhaSi76 [oxTi177; pDC731; Pdes-2::mScarlet-I::PH::unc-54::3'UTR; cb-unc-119(+)]IV; arx-2 (dha237 [arx-2::AID])V* | This study | DKC1426 |
| *dhaSi339 [pDC1158; Pdes-2::MYR::KNL-1(SAAA RRASA)::mTagBFP::unc-54::3'UTR; cb-unc119(+)]II; dhaSi243 [pDC929; oxTi177; Pdes‑2::Lifeact::mKate2::tbb-2::3’UTR; cb-unc-119(+)]IV; dhaSi45 [oxTi365; pDC675; Pdes-2::mNeonGreen::PH::unc-54::3'UTR; cb-unc-119(+)]V* | This study | DKC1429 |
| *dhaSi145 [oxTi185; pDC793; Pdes‑2::TIR1::F2A::mTagBFP::AID::NLS::tbb-2::3'UTR; cb-unc-119(+)]I; dhaSi89 [pDC747; Punc-86::TIR-1::unc-54::3'UTR; cb-unc-119(+)]II; knl-1 (dha122 [AID:: mTagBFP::knl-1]); cyk-1 (dha238 [cyk-1::AID])III; dhaSi243 [pDC929; oxTi177; Pdes-2::Lifeact::mKate2::tbb-2::3’UTR; cb-unc-119(+)]IV* | This study | DKC1430 |
| *dhaSi145 [oxTi185; pDC793; Pdes‑2::TIR1::F2A::mTagBFP::AID::NLS::tbb-2::3'UTR; cb-unc-119(+)]I; dhaSi89 [pDC747; Punc-86::TIR-1::unc-54::3'UTR; cb-unc-119(+)]II; knl-1 (dha122 [AID:: mTagBFP::knl-1])III; dhaSi76 [oxTi177; pDC731; Pdes-2::mScarlet-I::PH::unc-54::3'UTR; cb-unc-119(+)]IV; arx-2 (dha237 [arx-2::AID])V* | This study | DKC1441 |
| *dhaSi145 [oxTi185; pDC793; Pdes-2::TIR1::F2A::mTagBFP::AID::NLS::tbb-2::3’UTR; cb-unc-119(+)]I; dhaSi89 [pDC747; Punc-86::TIR-1::unc-54::3’UTR; cb-unc-119(+)]II; knl-1(dha122 [AID::mTagBFP::knl-1])III; dhaSi53 [oxTi177; pDC675; Pdes-2::mNeonGreen::PH::unc-54::3'UTR; cb-unc-119(+)]IV; dhaSi272 [oxTi365; pDC1122; Pser2-prom3::dma-1::mKate2::unc-54::3'UTR; cb-unc-119(+)]V* | This study | DKC1478 |
| *dhaSi145 [oxTi185; pDC793; Pdes-2::TIR1::F2A::mTagBFP::AID::NLS::tbb-2::3’UTR; cb-unc-119(+)]I; dhaSi89 [pDC747; Punc-86::TIR-1::unc-54::3’UTR; cb-unc-119(+)]II; dhaSi53 [oxTi177; pDC675; Pdes-2::mNeonGreen::PH::unc-54::3'UTR; cb-unc-119(+)]IV; dhaSi272 [oxTi365; pDC1122; Pser2-prom3::dma-1::mKate2::unc-54::3'UTR; cb-unc-119(+)]V* | This study | DKC1479 |
| *dhaSi207 [pDC1020; Punc-86::splitGFP1-10::tbb-2::3'UTR; cb‑unc‑119(+)]II; knl-1(dha130 [HA::7XGFP11::knl-1])III; dhaSi347 [oxTi177; pDC1278; Pdes-2::mScarlet-I::unc-54::3'UTR; cb-unc-119(+)]IV* | This study | DKC1482 |
| *dhaSi207 [pDC1020; Punc-86::splitGFP1-10::tbb-2::3'UTR; cb‑unc‑119(+)]II; dhaSi347 [oxTi177; pDC1278; Pdes-2::mScarlet-I::unc-54::3'UTR; cb-unc-119(+)]IV* | This study | DKC1483 |
| *dhaSi349 [pDC1216; Pdes-2::KNL-1::mTagBFP::unc-54::3'UTR; cb-unc119(+)]II; dhaSi243 [pDC929; oxTi177; Pdes‑2::Lifeact::mKate2::tbb-2::3’UTR; cb-unc-119(+)]IV; dhaSi45 [oxTi365; pDC675; Pdes-2::mNeonGreen::PH::unc-54::3'UTR; cb-unc-119(+)]V* | This study | DKC1504 |
| *dhaSi145 [oxTi185; pDC793; Pdes‑2::TIR1::F2A::mTagBFP::AID::NLS::tbb-2::3'UTR; cb-unc-119(+)]I; dhaSi89 [pDC747; Punc-86::TIR-1::unc-54::3'UTR; cb-unc-119(+)]II; knl-1 (dha122 [AID:: mTagBFP::knl-1]); cyk-1 (dha230 [cyk-1::AID])III; dhaSi243 [pDC929; oxTi177; Pdes‑2::Lifeact::mKate2::tbb-2::3’UTR; cb-unc-119(+)]IV; dhaSi45 [oxTi365; pDC675; Pdes-2::mNeonGreen::PH::unc-54::3'UTR; cb-unc-119(+)]V* | This study | DKC1518 |
